# Genome-wide DNA methylation patterns in Autism Spectrum Disorder and mitochondrial function

**DOI:** 10.1101/310748

**Authors:** Sofia Stathopoulos, Renaud Gaujoux, Colleen O’Ryan

**Affiliations:** Department of Molecular and Cell Biology, University of Cape Town, Cape Town, South Africa.; Cytoreason LTD.

**Keywords:** DNA methylation, epigenome, ASD, autism, mitochondria, metabolism, protein ubiquitination, gene co-expression modules

## Abstract

Autism Spectrum Disorder (ASD) is a neurodevelopmental disorder characterised by phenotypic heterogeneity and overlapping co-morbidities. The genetic architecture of ASD is complex, with 100’s of risk genes cumulatively contributing to the aetiology of ASD. Epigenetic mechanisms, particularly DNA methylation, have been associated with ASD. The vast majority of ASD molecular research has focused on Northern European populations, with a paucity of data from Africa. This study examines genome-wide DNA methylation patterns in a novel cohort of South African children with ASD and matched, unrelated controls. We performed a whole-genome DNA methylation screen using the Illumina 450K Human Methylation Array. We identify differentially methylated loci associated with ASD across 898 genes (p-value ≤ 0.05). Using a pathway analysis framework, we find nine enriched canonical pathways implicating 32 of the significant genes in our ASD cohort. These pathways converge on two crucial biological processes: mitochondrial metabolism and protein ubiquitination, both hallmarks of mitochondrial function. The involvement of mitochondrial function in ASD aetiology is in line with the recently reported transcriptomic dysregulation associated with the disorder. The differentially methylated genes in our cohort overlap with the gene co-expression modules identified in brain tissue from five major neurological disorders, including ASD. We find significant enrichment of three gene modules, two of which are classified as Mitochondrial and were significantly downregulated in ASD brains. Furthermore, we find significant overlap between differentially methylated and differentially expressed genes from our dataset with a RNA seq dataset from ASD brain tissue. This overlap is particularly significant across the Occipital brain region (padj= 0.0002) which has known association to ASD. Our differential methylation data recapitulate the expression differences of genes and co-expression module functions observed in ASD brain tissue which is consistent with a central role for DNA-methylation leading to mitochondrial dysfunction in the aetiology of ASD.

## Background

It is forecast that by 2050 (1) the African continent will be home to 40% of the world’s children, of whom 10–20 million are predicted to have Autism Spectrum Disorder (ASD). This prediction is based on the reported ASD incidence estimated in northern-hemisphere countries (2). There is a paucity of ASD information with little or no information available pertaining to epidemiology, diagnosis, intervention and particularly molecular genetics research from African populations (3). Currently there are only two publications reporting the genetic component of ASD in Sub-Saharan Africa (4), which highlights the importance of characterising the molecular basis of ASD in an African population. This knowledge gap is hamstrung by the ability to complete ADS genetic research in Africa, including South Africa, which is limited by the absence of ASD biological databanks with associated ASD phenotypes for African populations. This is in stark contrast to the European and North American ASD consortiums available to researchers working on those populations (4). South Africa has a unique population demography, and molecular investigations of a well-characterized ASD cohort can make a meaningful contribution to understanding the aetiology of this complex disorder in an understudied population(3,4).

ASD is a neurodevelopmental disorder characterised by deficits in social communication, restricted interests and repetitive behaviours (5). The genetic architecture of ASD is heterogeneous, with only 10% of ASD attributable to known genetic causes and 49% of ASD liability being quantitatively associated with common genetic variation (6). Although ASD is highly heritable, molecular investigations have yet to identify a consistent association of ASD with a biological marker (7). This reflects the complexity of the ASD phenotype, which is varied and overlaps with other co-morbidities (8). Although ASD is traditionally considered a brain disorder, a recent paradigm shift has emerged which considers ASD as a systems-level disorder (7,9), because of its numerous co-morbidities and the involvement of pleiotropic pathways in its aetiology. In this systems-level view, ASD is underpinned by a combination of overlapping biological networks that are dysregulated and affect multiple systems. The idea of overlapping biological networks being affected in ASD is reflected in the transcriptome, where post-mortem brain tissue of ASD individuals reveal distinct patterns of gene expression across a number of biological pathways (10–12). A recent study (9) using transcriptomic profiling coupled with GWAS data from five major psychiatric disorders including ASD, found shared pathways of transcriptional dysregulation, as well as disease-specific signatures of gene expression. The authors evaluated the genetic component of ASD using GWAS data in a subset of gene co-expression modules and raised the question of the contribution of epigenetic factors regulating gene expression, given that genetic variation accounted for a fraction of the transcriptional dysregulation observed in psychiatric disorders. Of the disease-specific transcriptional dysregulation signatures associated with the psychiatric disorders, specific gene co-expression modules involved in neuronal and mitochondrial function were downregulated in ASD, linking ATP energetics to synaptic transmission in psychiatric disorders (9).

Mitochondrial homeostasis is a fundamental physiological process and its dysfunction affects all tissues. Congenital mitochondrial diseases are often lethal or have severe consequences (13,14). There are more than 1500 gene products are involved in mitochondrial function (14) and expression changes of any of these genes can result in a range of mitochondrial dysregulation phenotypes. A 50 fold increase in mitochondrial disease in children with ASD (15), altered oxidative stress response in ASD lymphoblastoid cell lines (Rose et al. (16) and higher DNA damage, DNA hypomethylation and oxidised protein levels(17) have been reported. Given the high energetic demand of the brain, mitochondrial dysfunction has also been linked to other neurologic disorders (18). For example, multiple lines of evidence suggest that mitochondrial dysfunction plays a pivotal role in Parkinson’s Disease (PD) via mitochondrial respiration defects, aberrant programmed cell-death and mitophagy (mitochondrial-specific type of autophagy), abnormal mitochondrial fission, fusion and biogenesis (19,20).

The nuclear genome can epigenetically modulate mitochondrial function via histone modification, DNA methylation or metabolic signals from the mitochondria (21). It is accepted that the epigenome represents the interface between genes and the environment (22) with epigenetic mechanisms driving temporal and spatial gene regulation during cell differentiation and neurodevelopment (7,23,24). DNA methylation has increasingly been associated with complex behavioural phenotypes (25,26). For example, an epigenome-wide ASD monozygotic twin study revealed multiple differentially methylated genes associated with the disorder (27) and DNA methylation has been correlated with neuroimaging in ADHD-discordant monozygotic twins (28).

The enrichment of genetic ASD variants partly accounts for transcriptomic regulation in brain, with the role of epigenetic and environmental factors not yet fully characterised (9). This study aims to assess the contribution of DNA methylation in the differential gene expression observed in ASD. We identify differentially methylated genes associated with ASD a cohort of South African children using a whole-epigenome approach, and analyse these genes within a pathway analysis framework. The differentially methylated genes associated with ASD converge in pleiotropic pathways that result in mitochondrial dysregulation which implicate mitochondrial metabolic function and programmed cell-death via protein ubiquitination. Our data recapitulate the expression differences of genes and coexpression module functions observed in ASD brain tissue, supporting a central role of epigenetically-driven mitochondrial dysfunction in the aetiology of ASD.

## Results and Discussion

### ASD study cohort and phenotyping

There is a large body of knowledge regarding genetic variation and disease associations from Northern Hemisphere populations, with African populations being understudied (29). This bias is also observed in ASD genetic research (4) which is exacerbated the by the lack of a centralised ASD biobank for African populations. Therefore, the first step of our study was to build an ASD cohort of South African children with age- and gender-matched controls. All participants were phenotypically characterised using the autism assessment tool ADOS-2. Since autism is a spectrum disorder, its defining traits are quantitative and are present along a continuum which overlaps with typically developing individuals (30), thus all the control participants were also phenotypically assessed. We administered ADOS-2 assessments to 127 participants (72 case and 55 control), and found that 13.6% of the children in the case group, who had a prior independent diagnosis of ASD, did not meet ADOS-2 criteria for ASD. Furthermore, 18.2% of the children in the control group, were excluded because of the presence of defining autistic traits, as defined by the ADOS-2 criteria. This dropout rate highlights the importance of phenotyping controls, a procedure which is not common-practice in ASD studies, and allowed us to narrow the heterogeneity in our control cohort before proceeding to the molecular analyses.

### Differentially methylated genes in ASD

To examine the epigenetic component of ASD, we selected 48 study participants from our South African cohort (32 case and 16 control) to investigate differential DNA methylation using the Illumina 450K Human Methylation beadchip assay. We applied stringent quality control filtering criteria to our data, retaining 408604 CpG probes for subsequent analyses. To ensure that we were working with homogeneous epithelial cells samples, we tested methylation signals for the potential contamination from blood cells, which was confirmed to be absent. We tested for the covariates ethnicity and age, to confirm that these were not contributing to the observed differential methylation, and corrected for potential batch effects. We identified differentially methylated CpG sites across 898 genes between case and control, of which 39 were highly significant (p≤0.005, Figure 1, Table S1). Differentially methylated genes were distributed across all chromosomes, including the sex chromosomes (Figure 1). The presence of numerous differentially methylated genes on the Y chromosome is consistent with the gender bias observed in ASD (2). Of the 898 genes differentially methylated in our cohort, 34 have previous associations with ASD (from the human SFARI gene database, https://gene.sfari.org/database/human-gene/, Table S2) including two genes that are among the top differentially methylated genes in our cohort (p≤0.005): GLRA2, a ligand-gated channel mediating a chloride-dependent inhibitory neurotransmission (31), and AHI1, a gene involved in both cerebellar and cortical development in humans (32). In support of a role for epigenetic mechanisms in ASD aetiology, we also found two differentially methylated genes that are directly involved in DNA methylation and that had been previously associated to ASD: SETD5 and MTR. SETD5 is a methyltransferase within the 3p25 microdeletion; mutations of SETD5 have been identified in several studies of ASD probands, leading to the classification of this gene as a strong ASD candidate gene (33) (Table S2). MTR encodes the enzyme 5-methyltetrahydrofolate-homocysteine methyltransferase which catalyses the final step in methionine biosynthesis. MTR mRNA levels in postmortem human cortex were found to be significantly lower in autistic individuals than in controls, especially at younger ages, comparable to the age-range of our South African ASD cohort (34).

**Figure 1.**
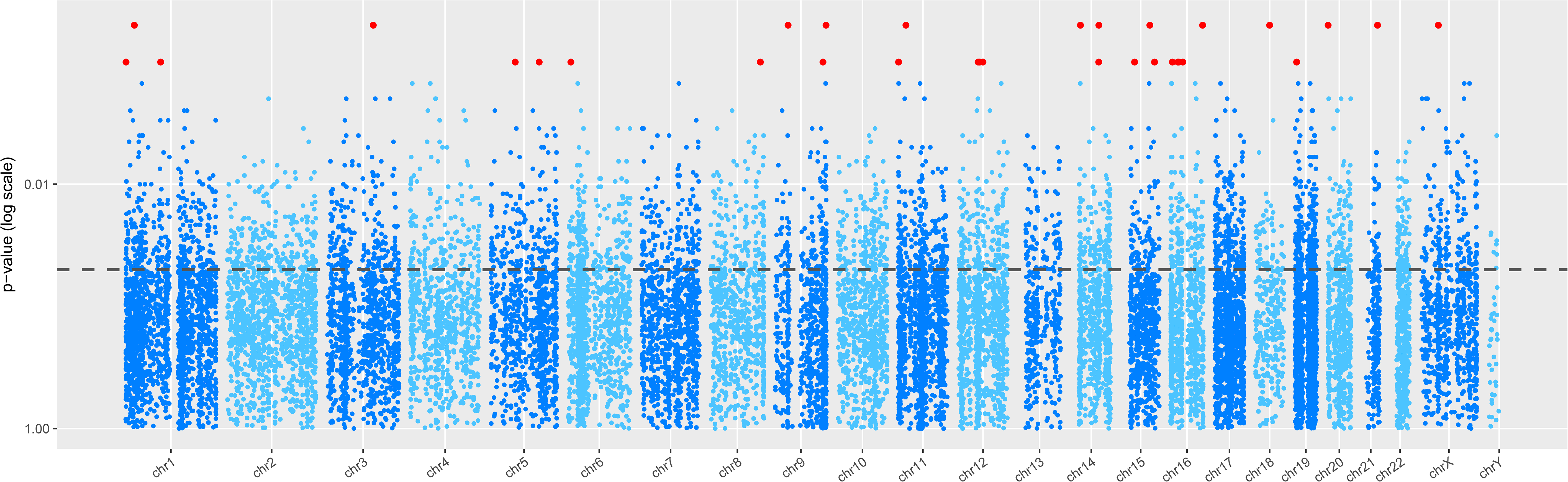
Differentially Methylated genes in a South African ASD cohort. Manhattan plot depicting genome distribution of differentially methylated genes in ASD. The dotted line denotes p-value = 0.05; highly significant genes (p≤0.005) are indicated in red.

### Canonical pathways implicated in ASD: Mitochondrial energetics

The limitations of single-gene analysis are apparent when examining the complicated genetic interplay typical of complex neurological disorders (35). Indeed, a number of transcriptomic studies have proposed that common pleiotropic pathways underpin not only ASD, but also a number of other neuropsychiatric disorders (9,36). Therefore, we analysed our data using a framework of pathway analysis, rather than a single-gene approach, to investigate the aetiology of ASD in our cohort.

We analysed the 898 genes with differentially methylated probes associated with ASD, in a canonical pathway framework. We identified nine enriched canonical pathways in our dataset (Table 1), which included 32 genes. Eight of the nine enriched canonical pathways identified in our dataset were metabolic pathways, with the majority of these occurring in the mitochondria (Table 1). Using IPA, we examined the overlap of the significant genes in our dataset with all the canonical pathways and functions associated with mitochondrial function and biogenesis. This revealed additional differentially methylated ASD genes in both the Mitochondrial Dysfunction canonical pathway and the Mitochondrial Biogenesis toxicity list (Table S3), which support the observed enrichment of mitochondrial functions in our dataset and a central role for mitochondrial homeostasis in ASD (37).

**Table 1.** Enriched Canonical Pathways and Associated Genes

| Ingenuity Canonical Pathways | -log(p-value) | Cellular location | Genes |
| --- | --- | --- | --- |
| Methylmalonyl Pathway | 2.06 | Mitochondria | MUT,PCCB |
| 2-oxobutanoate Degradation I | 1.85 | Mitochondria | MUT,PCCB |
| Glutaryl-CoA Degradation | 1.72 | Mitochondria | L3HYPDH,HACD2,EHHADH |
| Triacylglycerol Biosynthesis | 1.56 | Endoplasmatic Reticulum | LPGAT1,LPCAT3,PLPP5,MOGAT1,AGPAT1 |
| *Protein Ubiquitination Pathway | 1.51 | Cytoplasm | HSPA2,DNAJC13,PSMB5,DNAJC8,PSMC2,USP51,DNAJ C28,USP31,UBB,UBE2C,USP49,MDM2,PSMD10,SMURF 1,RBX1,DNAJC5G,PSMA5 |
| Acyl Carrier Protein Metabolism | 1.41 | Mitochondria | AASDHPPT |
| Acetyl-CoA Biosynthesis III (from Citrate) | 1.41 | Cytoplasm | ACLY |
| Lipoate Salvage and Modification | 1.41 | Mitochondria | LIPT1 |
| Superpathway of Methionine Degradation | 1.33 | Mitochondria | MTR,MUT,PCCB,PRMT1 |

\*Non-metabolic pathways

When synthesizing these seemingly disparate enriched metabolic canonical pathways within a framework of integrated metabolism, we observed that six of these pathways were directly linked to ATP production in the mitochondria (Figure 2). The Lipoate Salvage and Modification pathway transfers a lipoate moiety via the action of Lipoyltransferase 1 gene, LIPT1. This results in a lipoylated protein, the lipoyl-domain of two crucial enzymes of the Tricarboxylic acid cycle (TCA), pyruvate dehydrogenase (PDH) and alpha ketoglutarate dehyrogenase (a-KGDH). Similarly, the Glutaryl-CoA Degradation, 2-Oxobutanoate Degradation, Methylmalonyl pathway, Superpathway of Methionine Degradation Acyl Carrier Protein Metabolism, and Acetyl-CoA Biosynthesis III (from Citrate) all directly affect the substrates of the TCA, which culminates in the production of ATP.

**Figure 2.**
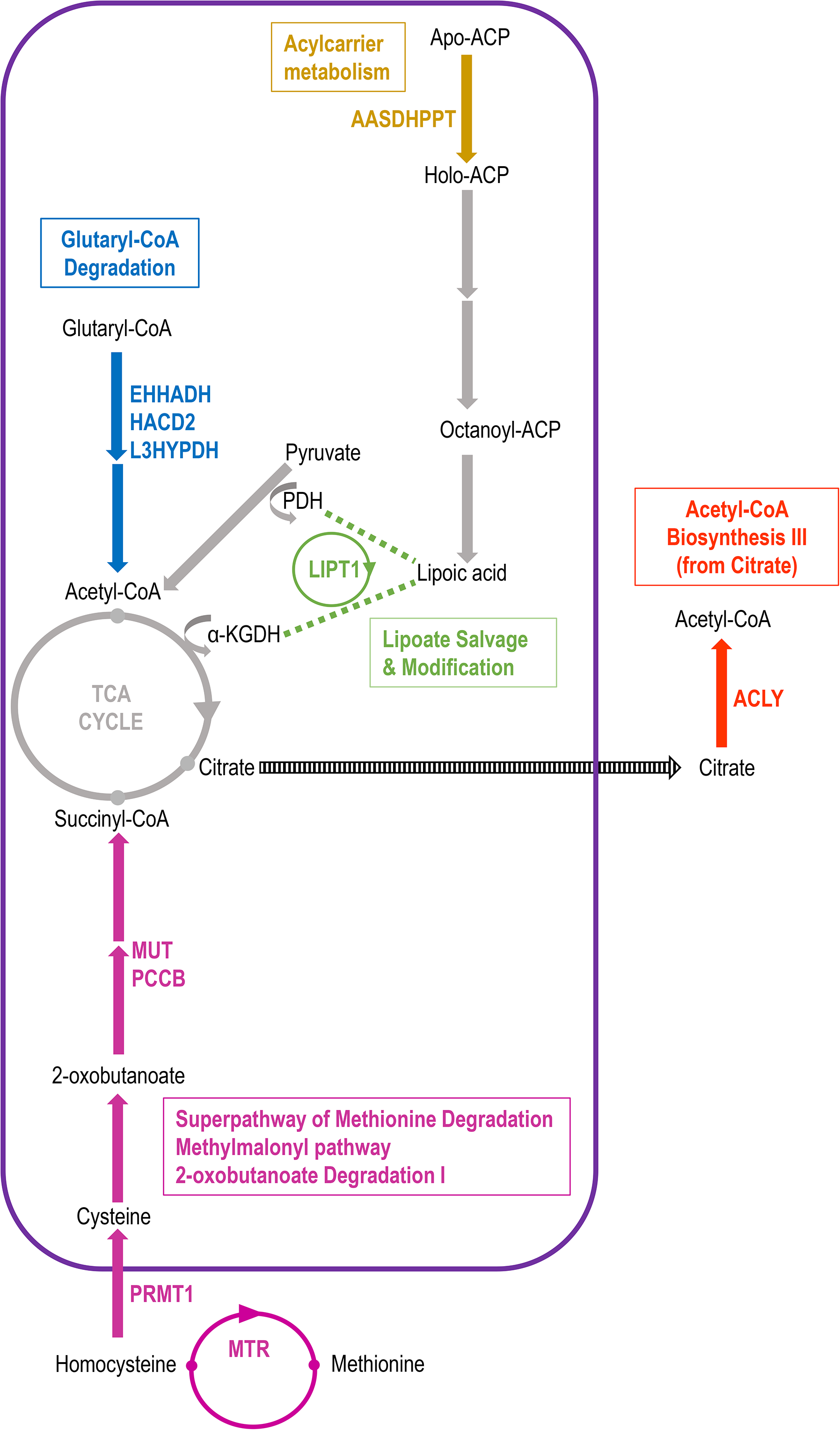
Mitochondrial pathways implicated in ASD aetiology. Differentially methylated genes in our dataset interact in multiple enriched canonical pathways (Table 2). When examining the enriched pathways in our ASD dataset, we find metabolic pathways which converge on the TCA cycle resulting in aberrant ATP production via the Electron Transport Chain, ETC, and the accumulation of Reactive Oxygen Species, ROS. The differentially methylated genes have a putative association with mitochondrial dysfunction in ASD. Differentially methylated genes are coded in colour corresponding to their respective canonical pathways which are indicated in colour-coded boxes. In these pathways, only relevant starting, intermediate and end-molecules are shown, where arrows can depict multiple reaction steps. Genes shown in colour are differentially methylated in our dataset resulting in changes in enzyme flux in their respective pathways.

In addition to the genes in the enriched canonical pathways, we found six differentially methylated genes associated with ASD that are part of the Mitochondrial Dysfunction canonical pathway (Figure 3: pathway, Table S3) that also affect ATP production and Reactive Oxygen Species (ROS). These include four genes from the NDUF family (NDUFA4, NDUFB2, NDUFB4, NDUFB6) which are part of complex I, one gene in complex III (UQCRC2) and one gene part of complex IV (COX7B) of the electron transport chain. When inspecting our top differentially methylated genes (p<0.005), we find that the most differentially methylated gene in our dataset, Stomatin Like 2 (STOML2), codes for a mitochondrial protein that regulates mitochondrial fusion in response to ROS damage. Importantly, the differential expressions of STOML2 and UQCRC2 have been validated in ASD brain tissue and contribute to two gene co-expression modules enriched in mitochondrial Gene Ontology pathways, which are downregulated in ASD (9).

**Figure 3.**
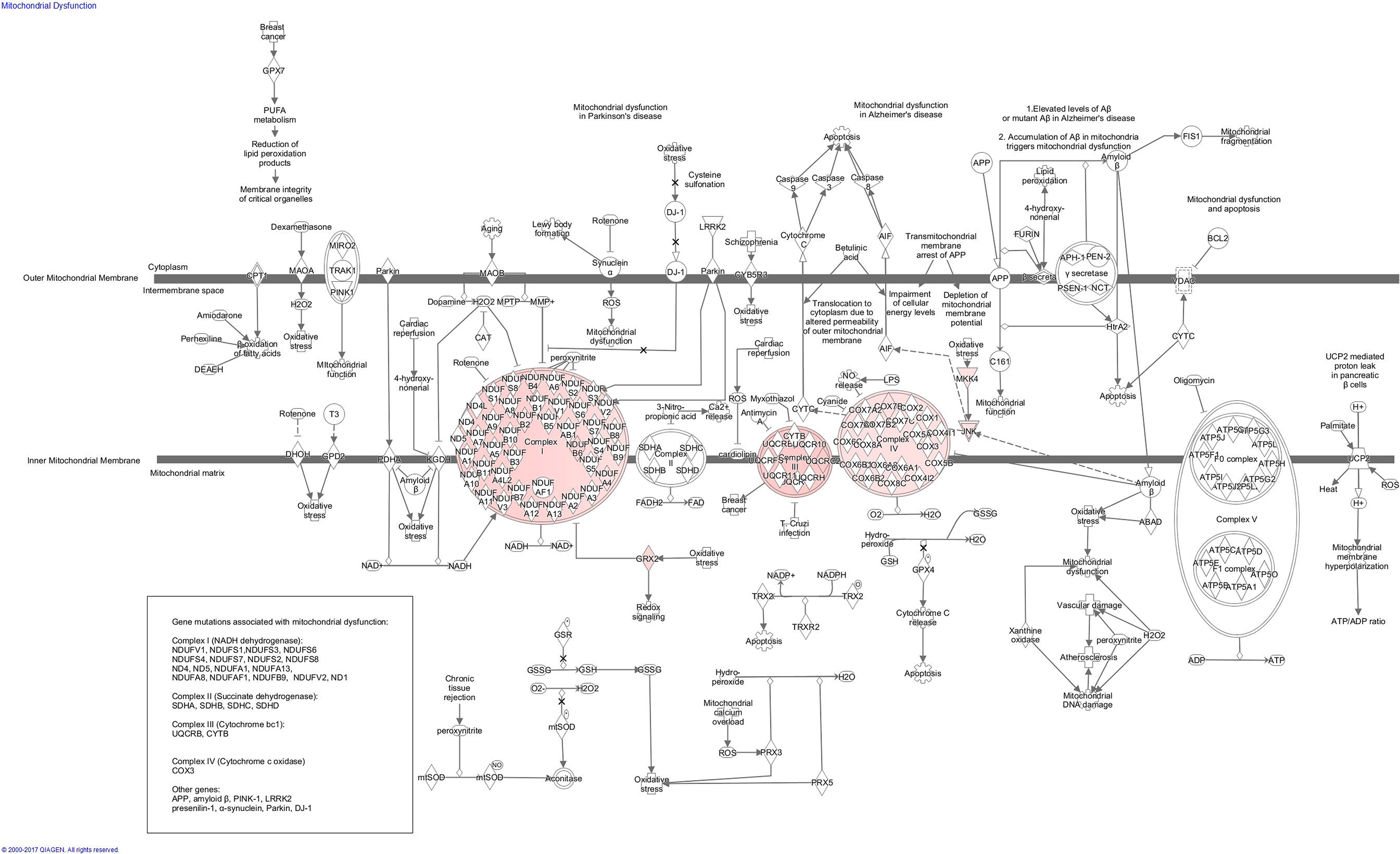
Differentially methylated genes in the Mitochondrial Dysfunction canonical pathway. Mitochondrial dysfunction canonical pathway (IPA; www.qiagen.com/ingenuity). We find eight genes that are differentially methylated within this canonical pathway in our South African ASD cohort: COX7B, GRX2, MAP2K4, NDUFA4, NDUFB2, NDUFB4, NDUFB6, UQCRC2. These genes are central to different steps of the Mitochondrial dysfunction pathway, which are coloured in light pink in this figure.

### Differentially methylated genes are enriched with co-expression modules linked to mitochondrial activity and ASD traits

To verify whether the results of our epigenomic screen were support by the gene expression differences observed in ASD brain tissue, we computed the enrichment our results in conjunction with available transcriptomic data (9). This recent study identified 13 gene co-expression modules that were robustly shared across ASD and four other psychiatric disorders in a cohort of 700 subjects of which there were 50 ASD subjects (9). The modules were defined from a co-expression network meta-analysis and were subsequently characterized according to their respective enrichment in GO categories and biological pathways, as well as their direct association with each psychiatric trait in the original cohort. Furthermore, the core expression pattern of each module was represented by a set of 20 top hub genes, as prioritized by their correlation with their respective module’s eigengene (9).

We assessed the concordance between the 13 modules and our results in three ways. First, we computed the overlap between the selected list of differently methylated genes from our data (898 genes) and the top 20 hub genes of each module (Fisher exact test). Secondly, we computed the overlap between the differently methylated genes from our data and all the genes in each module. Thirdly, we computed the enrichment of each module among the genes associated with the most differential methylation, using Gene Set Enrichment Analyisis (GSEA). Amongst the 13 modules, modules CD1, CD5, CD7 and CD10 showed a significant overlap with our ASD differentially methylated genes in at least one of the analyses (p-value ≤ 0.05, Figure 4). Of particular interest are modules CD5 and CD10, which were strongly associated with both mitochondrial activity and ASD traits, while module CD1 was associated with both ASD and neuron and synapse functions. All three modules (CD5, CD10 and CD1) were enriched in neuron-specific genes and were significantly down-regulated in ASD (9). The significant concordance between our epigenomic ASD signature and mitochondrial/neuronal enriched gene co-expression modules in ASD brains, is also supported by the association of mitochondrial gene-enriched modules with neuronal firing rate (37), linking ATP production to synaptic function in neuropsychiatric disorders.

**Figure 4.**
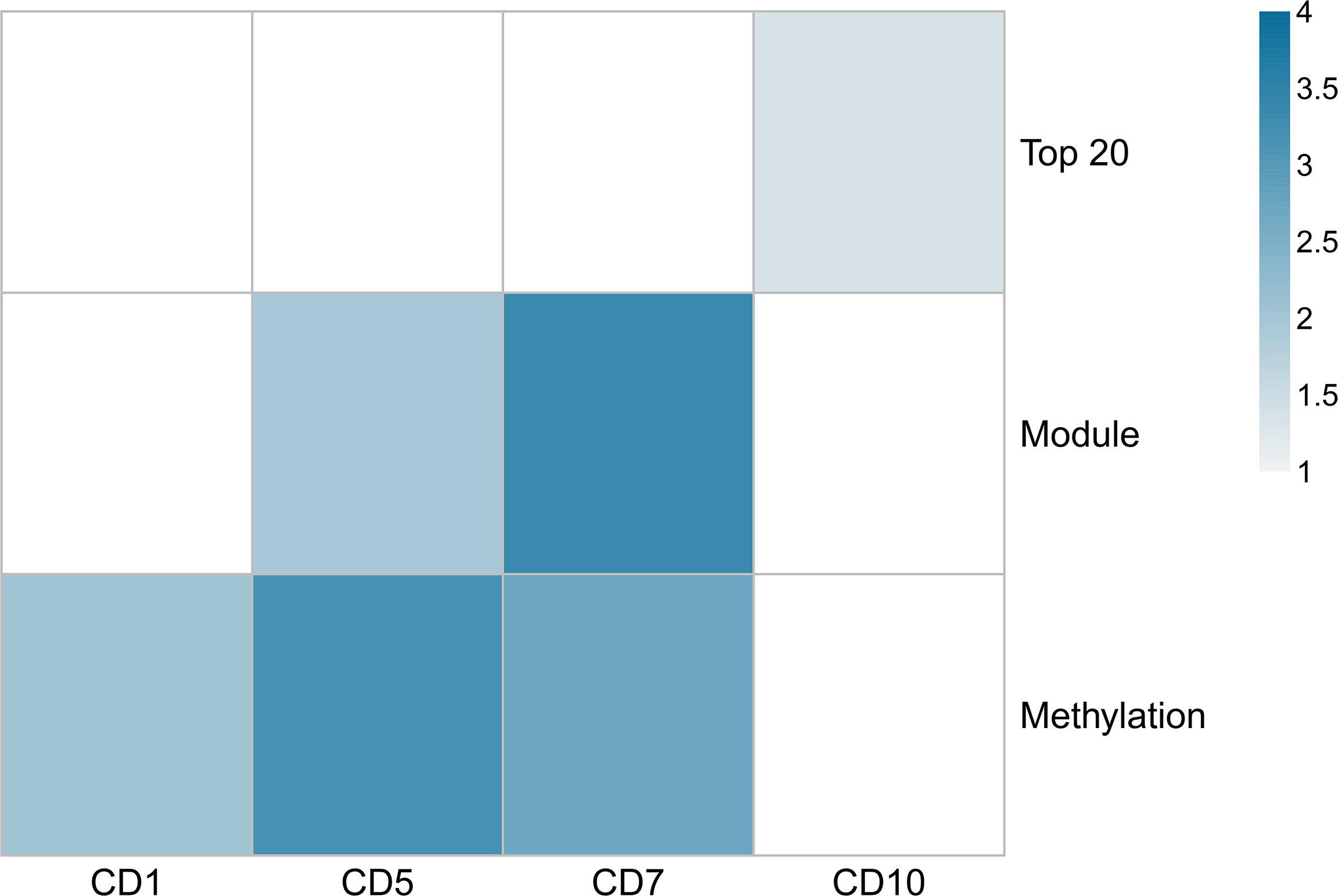
Differentialy methylated genes are enriched with co-expression modules linked to mitochondrial activity and ASD traits. Heatmap of −log10(p-value) of the comparisons of co-expression module overlap (‘Top 20’ and ‘Module’) and GSEA analysis (‘Methylation’) with our differentially methylated ASD genes. Only modules with p-value <= 0.05 are shown. The color indicates the strength of association, from weak (light blue) to strong (dark blue).

### Down-regulated gene expression in ASD occipital cortex is associated with differential methylation

Furthermore, we tested whether our differentially methylated ASD genes were associated with changes in gene expression in different brain regions. To do so, we used a published RNA-seq gene expression dataset (Synapse:syn4587609) that contains samples of post-mortem cortex brain of 24 ASD and 17 non-psychiatric subjects to obtain ASD gene expression signatures within four cortical regions (frontal, temporal, parietal and occipital lobes) (9). We used GSEA (38) to compute the enrichment of our list of differentially methylated genes (898 genes) in each brain region signature (ordered by moderated t-statistic). We found a significant, negative enrichment in the occipital brain region (Figure 5, Table S4) which suggests that the observed down-regulated gene expression in this brain region of ASD subjects may be linked to corresponding methylation differences. Interestingly, a recent study of brain imaging data from ASD and typically developing boys revealed a decrease of structural connectivity and resting-state brain activity in the occipital cortex of ASD brains, which is thought to be associated with the impaired social communication traits typical of ASD (39).

**Figure 5.**
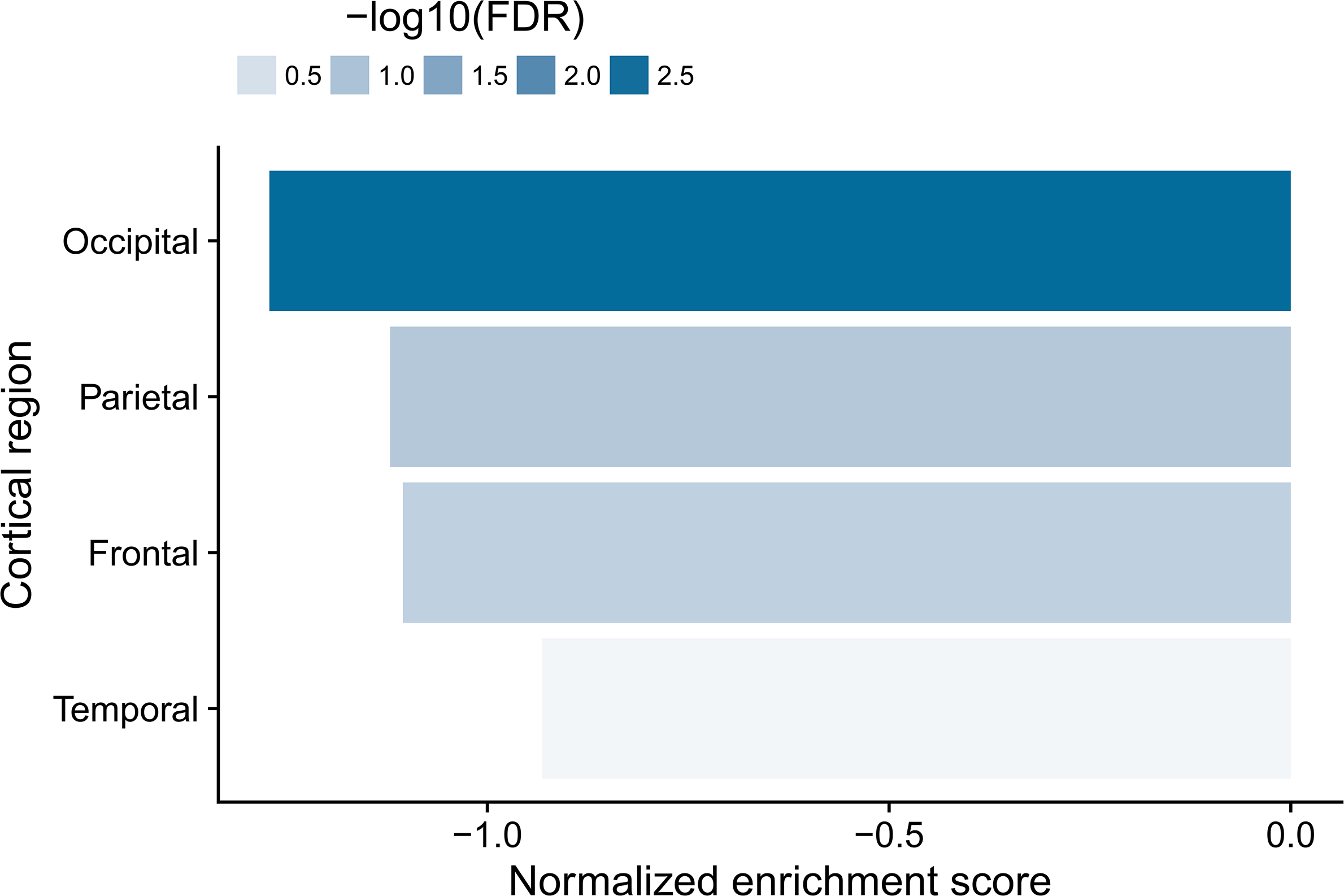
Down-regulated genes in the occipital region of ASD brain are enriched with differentially methylated genes. Normalized enrichment scores from GSEA in the ASD signature within each cortical region tested. Color indicates statistical significance from weak (light blue) to strong (dark blue).

Therefore, we propose that differential methylation of a combination of genes associated with ASD in our cohort (Table S1) results in the dysregulation of canonical pathways which culminate in mitochondrial dysfunction, leading to the many varied traits seen in ASD. Consistent with this hypothesis, the observed differential methylation of the genes in the enriched canonical pathways regulates the expression of their respective encoded enzymes (Figure 2), as is the significant overlap of our data with the transcriptomic data on ASD brain tissue. Furthermore, we propose that the flux of metabolites which ultimately affect the TCA cycle and ATP production, via the electron transport chain in the mitochondria, is perturbated in ASD. For example, the Lipoate Salvage and modification pathway enriched in our cohort, if dysregulated, would impact lipoic acid production. Lipoic acid is crucial for the TCA, with deletions of the gene impacting its biosynthesis or salvage resulting in early embryonic lethal phenotypes (40) or early onset fatal disease (41). Furthermore, among the differentially methylated genes in our ASD cohort that have a mitochondrial function is Glycine amidinotransferase (GATM), a gene encoding a mitochondrial enzyme involved in creatinine biosynthesis. Mutations of this gene lead to GATM deficiency, and were found to be associated with intellectual disability and behavioural disorders, including ASD, in a study of 27 patients (42).

Several studies have identified abnormal metabolic biomarkers in individuals with ASD (18). For example, a study by Filipek et al. (43) examined levels of serum biomarkers of oxidative phosphorylation in 100 children with ASD. The study revealed a combination of relatively low serum carnitine levels, slightly elevated lactate and significantly elevated alanine and ammonia levels, all suggestive of mild mitochondrial dysfunction in children with ASD (43). Furthermore, markers of abnormal ETC function have been identified in post-mortem brains of individuals with ASD, with decreased levels of complex I, complex III, complex IV and complex V proteins and impaired complex I and complex IV activity in temporal lobe of these brains (44). Noteworthy, most mitochondrial abnormalities were found in the brains of children under 10 years old, a similar age range as our cohort, suggesting a vulnerability of the developing brain (44).

Our mitochondrial hypothesis of epigenetic ASD aetiology is also consistent with the recent paradigm shift of considering ASD a systems-level disorder, rather than exclusively a brain disorder (7,45). Indeed, mitochondrial dysfunction has been proposed to explain the diverse symptoms associated with ASD (15,18,46). A recent systematic review and meta-analysis (15) based on a total of 18 publications examining children diagnosed with both ASD and mitochondrial disease, identified a 50-fold increase of mitochondrial disease in children with ASD, compared to the general population (15). Importantly, only 22% of these ASD individuals had a genetic cause underpinning mitochondrial disease, suggesting secondary mitochondrial dysfunction in the majority of cases. Mitochondrial dysfunction affects all tissues, particularly those with high energy demands like the central nervous system which depends on oxidative phosphorylation for successful neuronal development. It is therefore not surprising that defects in mitochondrial functioning have been associated with a range neurological disorders (47,48) including bipolar disorder (49), Alzheimer’s disease (50,51) and Parkinson’s disease (19,20).

### Canonical pathways implicated in ASD: Protein Ubiquitination and Mitophagy

Only one pathway of the nine enriched canonical pathways identified in our dataset was non-metabolic: the Protein Ubiquitination Pathway, with 17 differentially methylated genes (Table 1). These included SMURF1 (SMAD-specific E3 ubiquitin protein ligase 1) as well as four subunits of the proteasome (PSMA5, PSMB5, PSMC2 and PSMD10), all of which are crucial in the protein ubiquitination pathway, which leads to the degradation of damaged mitochondria (52). Mitophagy, the selective degradation of damaged mitochondria by autophagy, is a response to mitochondrial damage. Mitochondrial damage is “sensed” via complex protein interactions that include Pink1, Parkin and numerous translocase proteins of the mitochondrial outer membrane (TOM) complex, culminating in mitophagy via the ubiquitin – proteasome system (52–54). In our ASD cohort, we found three differentially methylated genes that are part of the Pink interaction network and 32 differential methylated genes in the Parkin (PRKN) interaction network, including STOML2, the top differentially methylated gene in our dataset (Table S1). Changes in TCA flux and oxidative phosphorylation, leading to the altered ATP and ROS production proposed to occur in ASD based on our results, could trigger molecular pathways that culminate in mitophagy which are similar to those observed in well-studied neurodegenerative diseases such as Parkinson’s Disease (54) and Alzheimer’s Disease (55).

Fifty additional differentially methylated genes in our dataset were associated with canonical pathways and networks of autophagy, apoptosis and other cellular functions central to cell death and survival (Table S5). It has been reported that a decrease in ATP leads directly to an increase in proteasome activity (56) with even mild oxidative stress increasing the activity of the ubiquitin – proteasome system (57). Supporting a role for ubiquitin – proteasome system in ASD, irrespective of the trigger for this system, is that five genes involved in ubiquitination (AHI1, CUL3, FBXO15, PSMD10 and UBA6) are also present in the SFARI database, and these genes are all differentially methylated in our dataset (Table S2). For example, CUL3 plays a role in polyubiquitination and the degradation of proteins, with rare mutations of this gene seen in individuals with ASD (58,59). Degradation of damaged mitochondria also occurs via caspase–dependent cell death (20), with caspase-3 proposed as a target to prevent the progression of Parkinson’s Disease (60). Caspases are a family of protease enzymes playing essential roles in programmed cell death and Caspase-2 is differentially methylated in our dataset (Table S5) and is involved in stress-induced cell death pathways and is implicated in neurodegenerative disorders like Alzheimer’s Disease, Huntington’s disease and temporal lobe epilepsy (61). Furthermore, Caspase-2 is inactivated by CSNK2A1, our third top differentially methylated gene associated with ASD. In addition, we found differentially methylated genes in the Apoptosis signalling and autophagy canonical pathways, as well as AHI1, which is linked to cell death (62). AHI1 is one our top differentially methylated genes (Tables S1 and S2) and it is associated with syndromic autism and Joubert syndrome (32); AHI1 is required for cerebellar and cortical development in humans. In addition, four more of our top differentially methylated genes (ASB15, EIF2B1, BLZF1 and EMP3) have functions that are part of the ubiquitin-proteasome system or are involved in autophagy (based on their respective IPA gene cards, www.ingenuity.com). Thus, the multiple differentially methylated genes identified in our data, the literature on neurodegenerative disorders and that reported on the SFARI data base, support the implication of the ubiquitin – proteasome system in ASD aetiology, in line with the well-studied role of this system mammalian brain formation (63).

### Conclusions

Our study is the first genetic study from Sub-Saharan Africa to examine a well-characterised ASD cohort using a whole-epigenome approach, and we identify novel differentially methylated genes associated with ASD. Our results highlight the need for ASD genetic research in under-studied populations (29). Our data supports a central role for DNA methylation in ASD aetiology which culminates in disrupted ATP homeostasis via mitochondrial dysfunction, suggesting a central role for epigenetic regulation in the recently reported transcriptomic ASD gene-expression signatures. Furthermore, our data suggests a role for aberrant protein degradation and mitophagy in ASD aetiology, consistent with what is observed in other well-described neurological conditions (20)(55). Given that there are a number of pre-existing therapeutic options already in use for mitochondrial dysfunction (18), our data identifies exciting new avenues for researching the treatment of ASD symptomology.

## Materials and Methods

### Cohort design

There is no ASD national database containing biological material and phenotypic profiles of individuals in South Africa. We recruited a cohort of children with ASD which were age- and gender-matched to typically developing controls. Given that methylation levels at some genes are associated with age (64), our age range was restricted to pre-pubertal children (6–12 years old) (65). Due to the higher genetic load and co-morbidities present in girls with ASD (66,67), we focused on boys only, in order to reduce the genetic heterogeneity in our cohort. Furthermore, only children with a prior independent diagnosis of ASD and without known co-morbidities were recruited into our case group; these children all attended special-needs Autism schools in the Western Cape, South Africa. We used the Autism Diagnostic Observation Schedule, ADOS-2, to characterise the ASD phenotype of each child. The level of ASD severity in our cohort ranges from Moderate to Severe according to the ADOS-2 scale. Our ASD cohort is therefore characterised by a range of phenotypes which are representative of the ASD spectrum. The control cohort consisted of genetically unrelated children with no diagnosis of ASD, ADHD, or any other developmental delays, attending ‘mainstream’ schools. In addition, the control cohort was screened for the complete absence of any of the traits that define ASD using ADOS-2. All ADOS assessments were done in English by a team of Research Reliable ADOS-2 administrators; recruited participants were instructed in English at their respective schools.

### Study participants and tissue collection

We screened 127 children, 72 cases and 55 controls, using ADOS-2 over a four-year period (2014–2017). Prior to recruiting, University of Cape Town ethical approval (FSREC076-2014) and Western Cape Government approval (20141002-37506) were obtained. Informed parental consent was obtained for each participant after we explained our study via oral and written presentations. Study participants did not receive a compensation and could withdraw from the study at any stage. Given the complex demographic history of South Africa, our study focused on the two demographic groups most represented at the recruiting sites: children of European and Mixed-Race ancestry. To limit the presence of the confounding factor of cellular heterogeneity, we used DNA extracted from buccal cells because these reflect DNA methylation of brain more accurately than blood cells (68,69). Buccal cells were collected using Epicentre Catch-All^™^ collection swabs. DNA was extracted using Proteinase K/ Salting out, followed by EtOH precipitation (70). DNA was re-suspended in sterile distilled water and quantified using Qubit 2.0 Fluorometer.

### Whole-epigenome methylation beadchip assay

DNA (500ng) from 48 children, 32 cases and 16 controls, was bisulphite converted using Zymo EZ DNA Methylation-Gold^™^ kit, as per manufacturer’s instructions. This sample size is similar to other methylome studies that were recently published (27,71), and reflects the semi-functional nature of the assay. Bisulphite-converted DNA samples were sent to the Agricultural Research Council core facility in Onderstepoort, South Africa for the beadchip assay. Genome-wide methylation was assessed using the Illumina Infinium HumanMethylation450 beadchip assay, which screens > 485 000 CpG sites across the entire genome at single-nucleotide resolution. The CpG target sites cover UTRs and the gene body of 99% of RefSeq genes, which include 96% of CpG islands and flanking regions. Arrays were scanned using Illumina HiscanSQ.

### Methylation beadchip analysis pipeline

All analyses were performed in R/Bioconductor (R Development Core Team 2011, (72)). Raw beadchip data were pre-processed and normalized using functions from the minfi, missMethyl and wateRmelon packages (73–75). Quality control of raw data was completed using the function wateRmelon::pfilter to flag low quality samples (over 1% sites with detection p-values > 0.05). The raw data for unflagged samples were then used to assess blood cell composition differences (minfi::estimateCellCounts) and normalized using SWAN (missMethyl::SWAN) (76). We then applied a sequence of filters to remove probes that had a detection p-value > 0.01 in at least one sample, probes that contained known SNPs (minfi::dropLociWithSnps) and probes that have been reported to be cross-reactive (77). After these filtering steps, the normalized data comprised 35 samples and 408 604 probes. All subsequent analyses were performed on M-values computed from these normalized data (with offset 100). Differences in methylation probe levels were obtained using the RUV-2 method (78), using Illumina 450k array negative controls and 0.5 FDR threshold for the first RUV fit. Differentially methylated probes (FDR <= 0.05) were annotated using Bioconductor annotation package FDb. InfiniumMethylation.hg19 which assigned methylation probes to the gene with the nearest transcription start site. A gene was called differentially methylated if it was assigned with at least one differentially methylated probe. In order to control for gene probe coverage differences, gene-level p-values were obtained using 2000 permutations of the sample labels, which generated null distributions that retain gene-specific coverage. P-values were computed by counting the proportion of times a gene obtained a statistic (maximum across assigned probes) in the permuted data, as extreme as the one observed on the original data.

### Pathway enrichment

In our case vs control analysis, we identified differentially methylated CpG sites across 898 genes. We used this gene list to identify the enriched pathways that differ between case and control using Qiagen’s Ingenuity Pathway Analysis, IPA (www.qiagen.com/ingenuity). All canonical pathways with a – log(p-value) > 1.3 are considered significantly enriched and we identified nine enriched pathways in our dataset, comprising a total of 32 differentially methylated genes. Of the 898 differentially methylated genes in our dataset, 39 genes have highly significant associations to ASD (p-value < 0.005). We use IPA to identify the enriched molecular networks associated with the differentially methylated genes in our cohort and known diseases and functions. We overlay our dataset with candidate canonical pathways which are interrelated with the core enriched canonical pathways in ASD. We find a significant enrichment of pathways consistent with mitochondrial dysregulation, implicating mitochondrial metabolic function and programmed cell-death via protein ubiquitination

### Enrichment analysis of gene co-expression modules and ASD cortical signatures

The co-expression modules and 20 top hub gene sets definitions were downloaded and extracted from Supplementary Table S2 from the recent paper reporting transcriptome profiling in brain tissue from individuals with ASD (n=50) and four other psychiatric disorders (n=700 individuals in total (9)). Gene symbols were standardized using the latest version of the Bioconductor Human annotation package (org.Hs.eg.db version 3.4.1). Overlap between the differentially methylated genes in our cohort and each module and module hub gene sets was assessed using a single-sided Fisher exact-test (alternative “greater”). Enrichment of each module in the differential methylation signature was computed using pre-ranked GSEA (Subramanian et al. 2005) via the fgsea package with 10000 permutations. Enrichment analysis in pan-cortical RNA-seq dataset was performed in a similar way, using the signatures defined in Supplementary Table S1 from Gandal et al. (2018) with standardized gene symbols and genes ordered by decreasing moderated t-test statistic in each region separately.

## List of abbreviations

AD: Alzheimer Disease
ADHD: Attention deficit hyperactivity disorder
ASD: Autism Spectrum Disorder
ADOS-2: Autism Diagnostic Observation Schedule-2
ATP: Adenosine triphosphate
ER: Endoplasmic Reticulum
ETC: Electron transport chain
GSEA: Gene Set Enrichment Analysis
GWAS: Genome Wide Association Study
IPA: Ingenuity pathway analysis
PD: Parkinson Disease
ROS: Reactive Oxygen Species
TCA: Tricarboxylic acid cycle

## Declarations

### Ethics approval and consent to participate

University of Cape Town ethical approval (FSREC076-2014) and Western Cape Government approval (20141002-37506) were obtained to conduct all aspects of this study. Informed parental consent was obtained for all study participants; study participants did not receive a compensation and could withdraw from the study at any stage.

### Availability of data and material

The datasets used and/or analysed during the current study are available from the corresponding author on reasonable request.

### Authors’ contributions

COR and SS conceptualised and designed the study, performed all experiments, analysed and interpreted the results and wrote the manuscript. RG implemented the bioinformatic pipeline for data analysis. All authors read and approved the final manuscript.

## Acknowledgements

We thank the families and the staff of all schools that participated in our research.

